# Sustained *Ranavirus* outbreak causes mass-mortality and morbidity in imperiled amphibians

**DOI:** 10.1101/2021.10.15.464511

**Authors:** Arik M. Hartmann, Max L. Maddox, Robert J. Ossiboff, Ana V. Longo

## Abstract

A persistent two-month long outbreak of *Ranavirus* in a natural community of amphibians contributed to a mass die-off of gopher frog tadpoles *(Lithobates capito)* and severe disease in striped newts *(Notophthalmus perstriatus)* in Florida. Ongoing mortality in *L. capito* and signs in *N. perstriatus* continued for five weeks after the first observation. Hemorrhagic disease and necrosis were diagnosed from pathological examination of *L. capito* tadpoles. We confirmed detection of a Frog Virus 3 (FV3)-like *Ranavirus* via quantitative PCR in all species. Our findings highlight the susceptibility of these species to *Rv* and the need for long-term disease surveillance during epizootics.

## Introduction

Emerging wildlife diseases are increasingly associated with amphibian mass mortalities and global amphibian declines (Rachowicz et al. 2006), and have led to heightened awareness and surveillance of amphibian pathogens. Iridoviruses in the genus *Ranavirus* (*Rv*) and the amphibian chytrid fungus *Batrachochytrium dendrobatidis* (*Bd*) are two emerging pathogens that are widely associated with amphibian mass mortality events (Miller et al. 2011; Fisher and Garner 2020). Outbreaks often result in high mortality of sensitive life stages or species, while tolerant species and life stages can serve as pathogen reservoirs (Gray et al. 2009; Schloegel et al. 2010). In the United States (US), pathogen-mediated mass mortalities and declines have been primarily recorded in larval amphibians in the northern and western regions of the country (Green et al. 2002). The Southeastern Coastal Plain of the US harbors the highest diversity of amphibians in North America (Noss et al. 2015), and although *Rv* and *Bd* have been detected in the region, reports of mortality events and their effects are lacking.

We report here the findings of a two-month long outbreak of *Rv* in a natural amphibian community using pathological examination of moribund tadpoles and confirmed pathogen presence via quantitative PCR (qPCR). We present the first report of *Rv*-induced mass mortality and morbidity in two Coastal Plain endemic amphibians: the gopher frog *(Lithobates capito)* and the striped newt *(Notophthalmus perstriatus).* Both species have histories of range-wide declines (Jensen and Richter 2005; Farmer et al. 2017), and are listed by the Florida Fish and Wildlife Conservation Commission (FWC) as species of greatest conservation need (FWC 2019).

We first observed mass-mortality of *L. capito* tadpoles on 23 January 2021 at One Shot Pond, in Ordway-Swisher Biological Station (OSBS), Putnam County, Florida. We observed ongoing die-offs during a second visit to the pond on 7 March 2021. One Shot Pond is a semi-permanent fishless wetland that provides important breeding habitat for 16 amphibian species, and is one of few wetlands at OSBS that supports both *L. capito* and *N. perstriatus* populations (LaClaire 1995; Johnson 2002).

During both surveys, we captured amphibians by dipnet or hand and stored them in individual plastic bags for processing. Each amphibian was examined to confirm species identification and detect gross symptoms of disease. For pathogen sampling, we swabbed the oral disc and vent of tadpoles following procedures established by (Gray et al. 2012) and swabbed caudates and post-metamorphic anurans following standard protocols for amphibian disease sampling (Hyatt et al. 2007). We collected 19 dead and moribund *L. capito* tadpoles for histopathological analysis and fixed them in 70% ethanol. We estimated over 500 *L. capito* tadpoles at Gosner stages 28-31 (Gosner 1960) had died between the two events, and most living tadpoles of the same age class showed symptoms of *Rv* infection, such as edema and hemorrhage (Fig. 1A-B). We did not observe mortality in *N. perstriatus*, however all newts exhibited erythema, hemorrhage, or necrosis (Fig. 1C-D). *L. capito* tadpoles at Gosner stages 23-25 (hatchlings) were abundant and did not show any clinical signs of infection, nor did any southern cricket frogs *(Acris gryllus)* around the pond perimeter.

**Figure 1.**
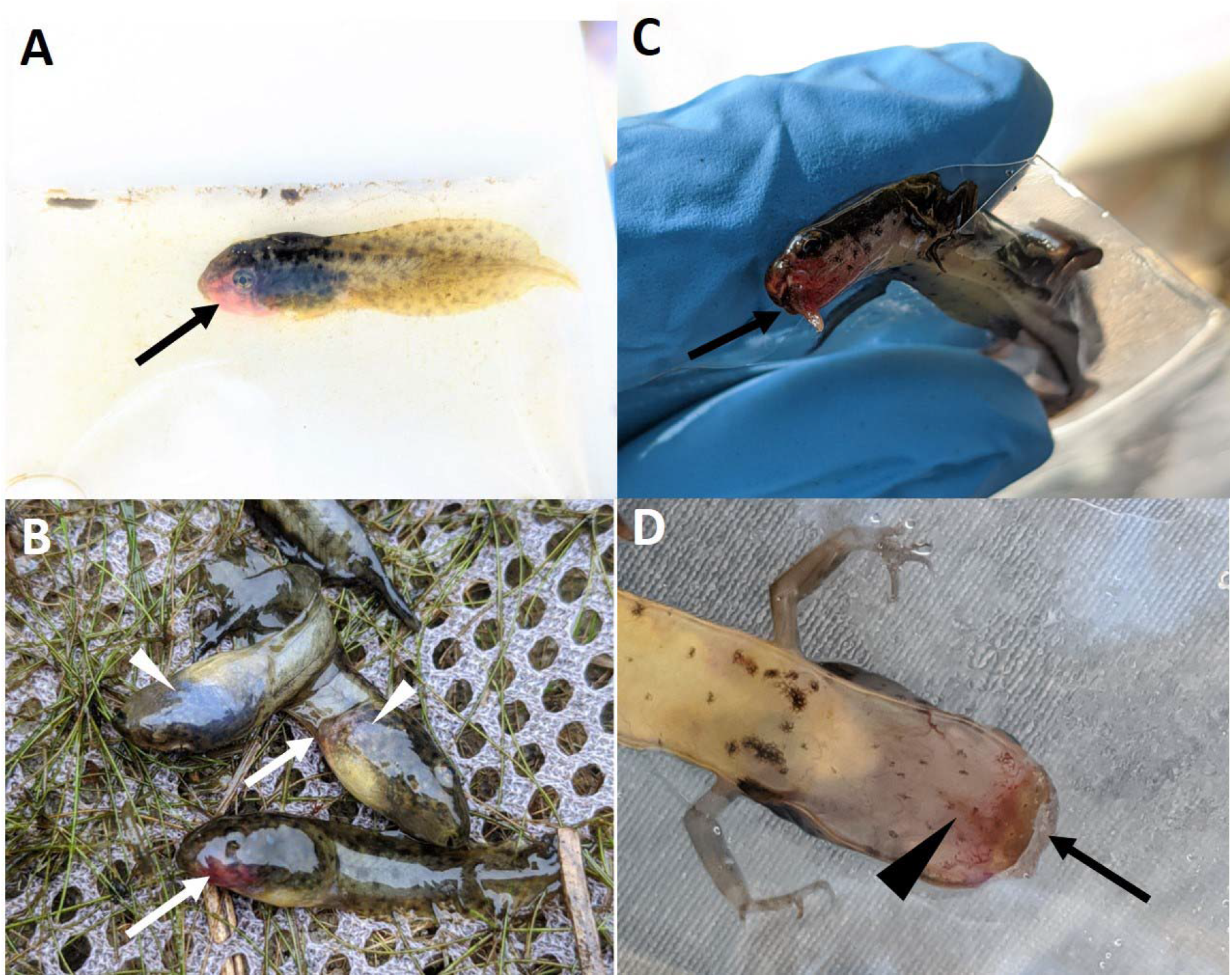
Gross symptoms of disease in *Ranavirus-infected* amphibians at One Shot Pond. (A-B) Moribund gopher frog (*Lithobates capito*) tadpoles showing hemorrhages (arrows) and discoloration (triangles). (C-D) Paedomorphic striped newts *(Notopthalmus perstriatus)* showing hemorrhage and necrosis (arrows), and erythema (triangle) of the mouth.

All 19 ethanol-fixed tadpoles were briefly decalcified in 0.5 M ethylenediamine tetraacetate acid (EDTA), pH 8.0 for ~24 h before sagittal sectioning and routine histopathologic processing and staining. Microscopic findings included necrosis of the genal glomeruli and interstitium (16/19 tadpoles, Fig. 2A), spleen (3/19 tadpoles, Fig. 2B), and liver (2/19 tadpoles) with cutaneous and subcutaneous hemorrhage (4/19 tadpoles) and vascular inflammation (2/19 tadpoles). In a subset of tadpoles (8/19), there were basophilic to amphophilic, cytoplasmic viral inclusion bodies present in hepatocytes (Fig. 2C). Swab samples were tested for FV3-like *Rv* and *Bd* using qPCR assays following protocols established by (Allender et al. 2013) and (Boyle et al. 2004), respectively. We mostly detected severe *Rv* infections in *L. capito* tadpoles, paedomorphic and recently metamorphosed *N. perstriatus,* and low to moderate infections in adult *A. gryllus* from both sampling events (Fig. 3). Species with disease signs had a higher proportion of individuals with intense *Rv* infections (>10^5^+). We detected *Bd* only in 10 *A. gryllus,* nine of which were also co-infected with *Rv* (Table 1).

**Figure 2.**
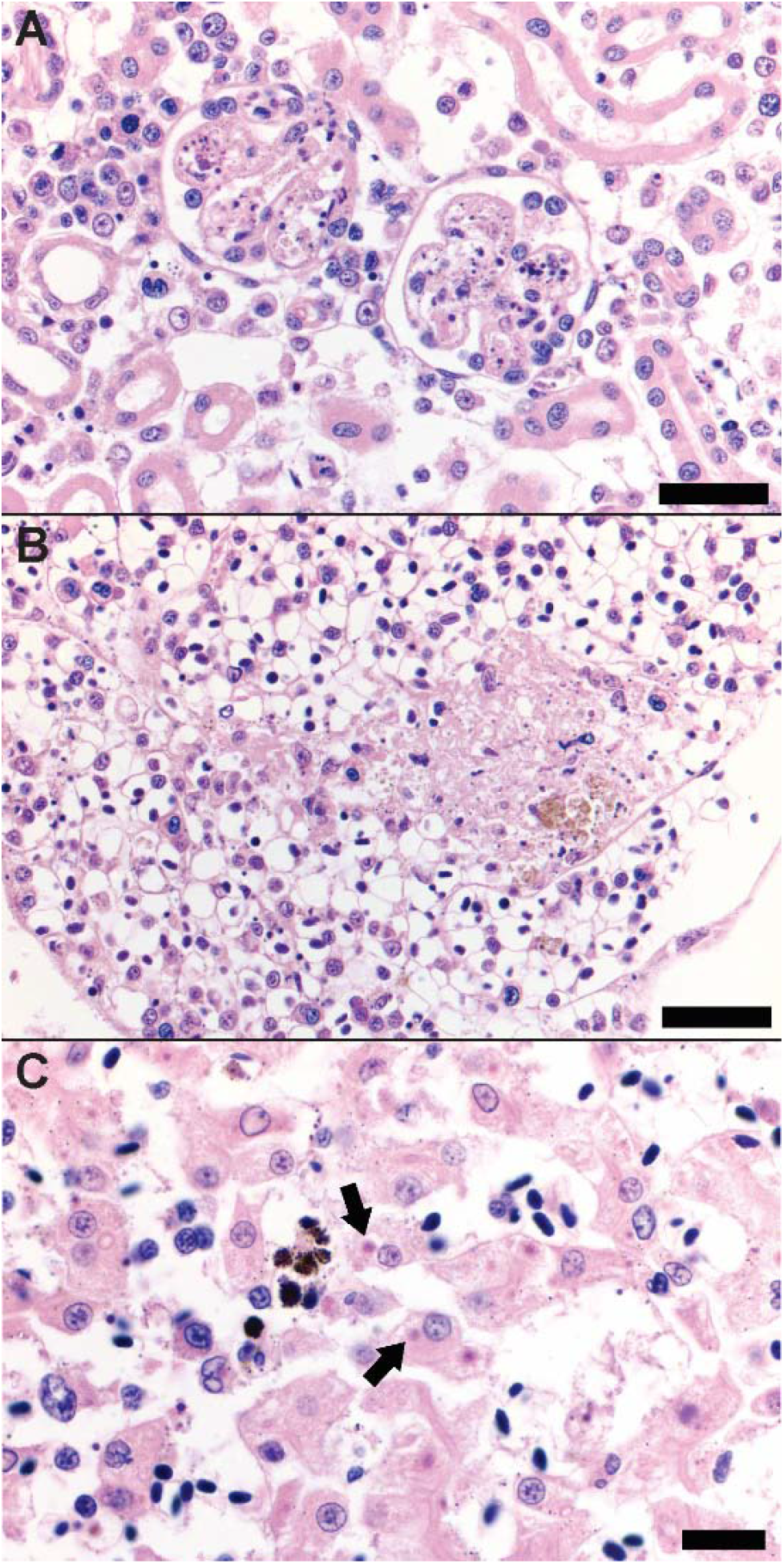
Histologic evidence of *Ranavirus* infection in gopher frogs *(Lithobates capito).* (A) Renal glomerular necrosis [Bar = 50 microns]. (B) Splenic necrosis [Bar = 50 microns]. (C) Cytoplasmic ranaviral inclusions highlighted by black arrows [Bar = 20 microns].

**Figure 3.**
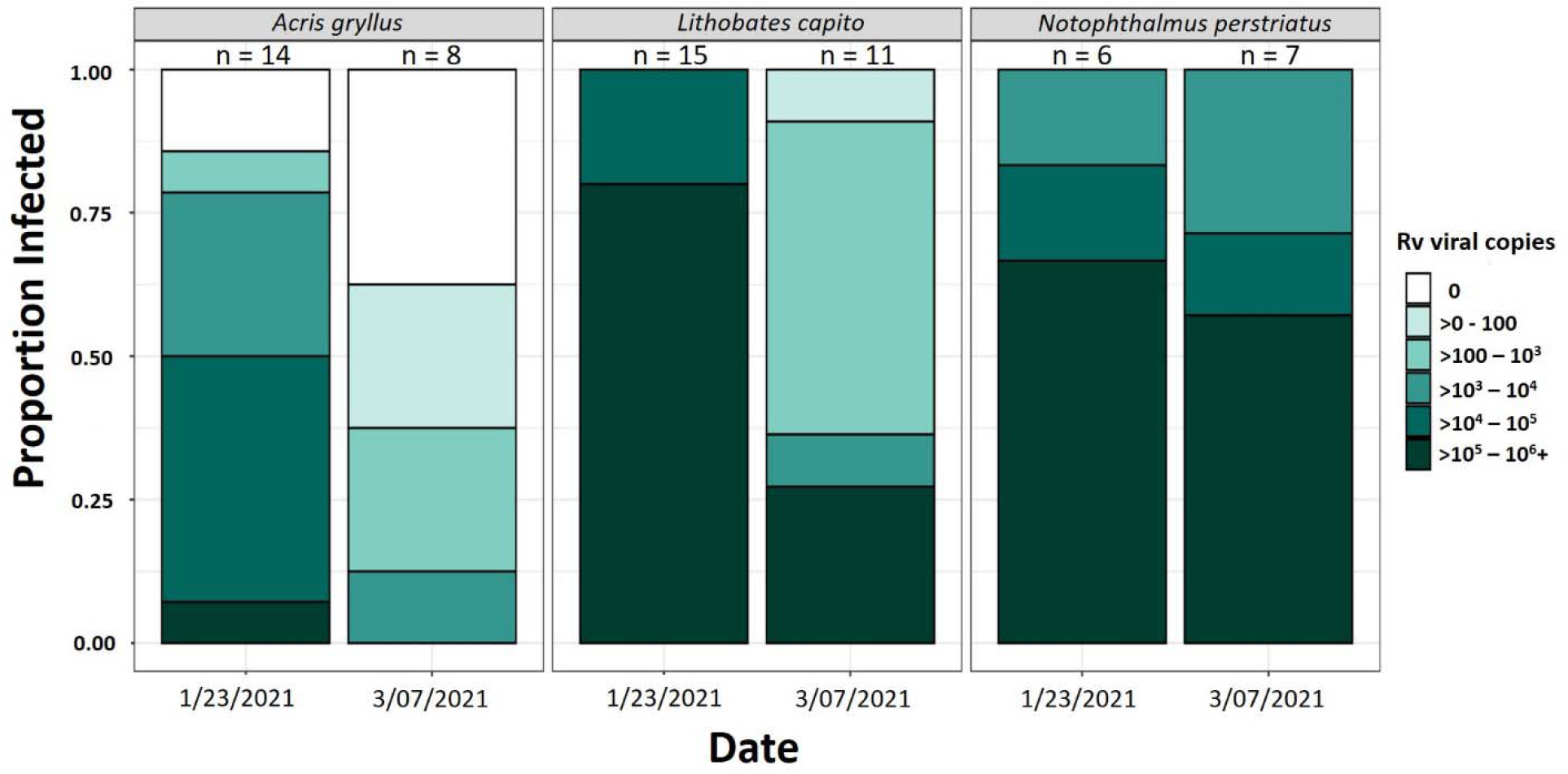
Prevalence and intensity of *Ranavirus* infections in three amphibian species during two sampling events at One Shot Pond at the Ordway-Swisher Biological Station.

**Table 1.** Species, life stage (A = adult, L = larva, P = paedomorph, MP = metamorphosing paedomorph) and prevalence of infection by *Rv* and *Bd* and the average *Rv* intensity combined from the two sampling events.

| Species | Life Stages Detected | Number sampled | *Rv*+ | (Prev.) | *Bd*+ | (Prev.) | *Bd*+/*Rv*+ | (Prev.) | Avg. *Rv* intensity (viral copies) |
| --- | --- | --- | --- | --- | --- | --- | --- | --- | --- |
| *Acris gryllus* | A | 22 | 17 | (77.2%) | 10 | (45.5%) | 9 | (40.9%) | 844.15 |
| *Lithobates capito* | L | 26 | 26 | (100%) | 0 | (0%) | 0 | (0%) | 2.3 x 10<sup>6</sup> |
| *Notophthalmus perstriatus* | P, MP | 13 | 13 | (100%) | 0 | (0%) | 0 | (0%) | 1.2 x 10<sup>6</sup> |
| **Total** |  | **61** | **54/61** | **(88.5%)** | **10/61** | **(16.4%)** | **9/61** | **(14.8%)** | **1.3 x 10<sup>6</sup>** |

*Rv* outbreaks can impact amphibian population dynamics by dampening recruitment, and pathogen persistence in the environment can negate recruitment entirely (Petranka et al. 2007), facilitating the local extinction of rare species (Earl et al. 2016). To our knowledge, reports of *Rv* outbreaks in natural populations of *L. capito* have not been published, but experimental infections of *L. capito* resulted in >90% mortality of tadpoles (Hoverman et al. 2011). Our observations show similar susceptibility under natural conditions, providing support for disease-related declines. Ongoing die-offs of tadpoles suggest that older cohorts may serve as vectors to younger cohorts through viral shedding, direct contact, or necrophagy (Harp and Petranka 2006; Peace et al. 2019).

As a multi-host pathogen of ectotherms, *Rv* outbreaks in amphibians can spread to the wider ectothermic community (Brenes et al. 2014). Many chelonians are susceptible to *Rv*, including federally protected gopher tortoises *(Gopherus polyphemus)* (Johnson et al. 2008; Cozad et al. 2020), and *Rv* outbreaks in chelonians have been attributed to pathogen spillover from sympatric amphibians (Brunner et al. 2015). *L. capito* are closely associated with *G. polyphemus* and are one nine amphibians known to cohabitate in tortoise burrows (Jackson and Milstrey 1989). At One Shot Pond there are >10 *G. polyphemus* burrows within 30 meters of the pond, and we have observed *L. capito* calling from burrow entrances. It is possible that *Rv* outbreaks can spill over to *G. polyphemus* and other ectothermic commensals by adult *L. capito* moving between ponds and burrows during breeding.

In contrast to *L. capito,* natural populations of *N. perstriatus* have not been extensively surveyed for disease, but *Rv* is a common pathogen of the closely related and sympatric eastern newt (*N. viridescens)* (Rothermel et al. 2016). *N. perstriatus* have experienced enigmatic declines and extirpations throughout their range in Florida and Georgia (Farmer et al. 2017), and repatriation efforts have been unsuccessful (Means et al. 2017). Experimental *Rv* exposure of captive reared *N. perstriatus* resulted in high mortality of aquatic and recently metamorphosed stages (Means et al. 2016). Our observations suggest that high pathogen pressure could result in *N. perstriatus* declines, and persistent disease may be inhibiting the recovery of the species. Our results provide evidence of disease-related risks in populations, a missing element that can strengthen the petition to list *N. perstriatus* under the Endangered Species Act (USFWS 2011; 2016).

In species with complex life histories, densities and life-stages fluctuate seasonally and can result in recurring epidemics and pathogen persistence through transmission between life stages (Brunner et al. 2004). *N. perstriatus* have complex life history strategies that include facultatively paedomorphic and triphasic developmental routes, and spend three of five life stages in water (Johnson 2002). These findings support our ongoing studies where we have found that paedomorphic life stages are more susceptible to *Rv* and experience higher disease burdens than other life stages (Hartmann et al. in preparation). We hypothesize that pathogen pressure in aquatic stages is density-dependent. Sustained *Rv* infections in metamorphosing newts may allow them to act as intraspecific reservoirs when they return to ponds as adults to breed. Long-term *Rv* persistence may select against the paedomorphic developmental route in *N. perstriatus*, which would have profound effects on population structure and annual recruitment as paedomorphic stages undergo accelerated maturation and reproduction (Dodd 1993).

Our findings also identify tolerant hosts that may act as reservoirs to more susceptible species (Brunner et al. 2004). *A. gryllus* are often the most abundant amphibian at ponds within OSBS (personal observation), occupy a variety of habitats, and can easily disperse between water bodies within OSBS (Dodd 1996). Because we did not find disease signs in *A. gryllus*, we hypothesize that high tolerance may allow *Rv* to persist in these hosts. Estimating dispersal rates for this species can help us predict pathogen spread across habitat types and amphibian assemblages.

Despite the diversity of amphibians in Florida and history of recent declines, reports of amphibian dieoffs in the state are rare and few have been published (Landsberg et al. 2013). Although both *Rv* and *Bd* have been detected, prior reports of mass mortality events in Florida have been attributed to Perkinsea parasites (Isidoro-Ayza et al. 2017). Here we provide the first report of a mass mortality event attributed to *Rv* in *L. capito.* The confirmation of *Rv* infection, resulting disease, and mass mortality pose major concerns for Florida’s amphibian and reptile populations, particularly specialist species with limited ranges. Both *L. capito* and *N. perstriatus* are habitat specialists in Florida threatened by habitat loss (Enge et al. 2014), and it is in the interest of state and federal wildlife agencies to further explore the implications that emerging pathogens have on management strategies. Future work must consider the role of emerging pathogens in past and continued amphibian declines. Current conservation plans must include pathogen mitigation strategies to ensure population survival and success of repatriation programs.

## Acknowledgements

We thank B. Folt (USGS) and L. Brendel (UF) for helping with sample collection, and the staff of the Histology Laboratory at University of Florida’s College of Veterinary Medicine for slide preparation. We also thank Andy Rappe and the staff at the Ordway-Swisher Biological Station for facilitating this work. Sampling was carried out under the permission of the Florida Fish and Wildlife Conservation Commission (FWC-LSSC-17-00031B) and the University of Florida’s Institutional Animal Care and Use Committee (#201810502). We thank members of the Longo Lab and S. Cassidy for their feedback on previous drafts of this manuscript.

## Notes

### Competing Interest Statement

The authors have declared no competing interest.

